# ENVIREM: An expanded set of bioclimatic and topographic variables increases flexibility and improves performance of ecological niche modeling

**DOI:** 10.1101/075200

**Authors:** Pascal O Title, Jordan B Bemmels

**Affiliations:** Department of Ecology and Evolutionary Biology, University of Michigan, Ann Arbor, Michigan 48109, USA

**Keywords:** bioclimatic variables, ecological niche model, Last Glacial Maximum, Maxent, predictor variable selection, species distribution modeling, WorldClim

## Abstract

Species distribution modeling is a valuable tool with many applications across ecology and evolutionary biology. The selection of biologically meaningful environmental variables that determine relative habitat suitability is a crucial aspect of the modeling pipeline. The 19 bioclimatic variables from WorldClim are frequently employed, primarily because they are easily accessible and available globally for past, present and future climate scenarios. Yet, the availability of relatively few other comparable environmental datasets potentially limits our ability to select appropriate variables that will most successfully characterize a species’ distribution. We identified a set of 16 climatic and two topographic variables in the literature, which we call the envirem dataset, many of which are likely to have direct relevance to ecological or physiological processes determining species distributions. We generated this set of variables at the same resolutions as WorldClim, for the present, mid-Holocene, and Last Glacial Maximum (LGM). For 20 North American vertebrate species, we then assessed whether including the envirem variables led to improved species distribution models compared to models using only the existing WorldClim variables. We found that including the ENVIREM dataset in the pool of variables to select from led to substantial improvements in niche modeling performance in 17 out of 20 species. We also show that, when comparing models constructed with different environmental variables, differences in projected distributions were often greater in the LGM than in the present. These variables are worth consideration in species distribution modeling applications, especially as many of the variables have direct links to processes important for species ecology. We provide these variables for download at multiple resolutions and for several time periods at envirem.github.io. Furthermore, we have written the ‘envirem’ R package to facilitate the generation of these variables from other input datasets.

## Introduction

The ability to model a species’ geographic distribution, given occurrence records and environmental information, is based on the assumption that abiotic factors directly or indirectly control species distributions (Austin 2002). Species distribution modeling (SDM) has led to a surge in research on topics such as species’ potential invasiveness (Thuiller et al. 2005), the impacts of climate change on species distributions (Thuiller 2004, Hijmans and Graham 2006, Morin and Thuiller 2009), the relative importance of various predictors in determining species range boundaries (Glor and Warren 2010), historical reconstructions of species distributions (Svenning et al. 2011), conservation applications such as the identification of suitable habitats for undiscovered populations or reintroductions (Martínez-Meyer et al. 2006), analysis of broad-scale patterns of species richness (Pineda and Lobo 2009), and spatially-explicit demographic simulations (Chan et al. 2011, He et al. 2013). The ability to conduct such analyses at increasingly broad taxonomic and spatial scales has largely been facilitated by successful efforts to digitize museum specimen records, georeference associated localities (Guralnick et al. 2006, Ellwood et al. 2015) and provide this information in a standardized format through easily accessible data portals (Constable et al. 2010, Wieczorek et al. 2012). While progress has been made in these efforts to make high quality occurrence records widely available (e.g., Global Biodiversity Information Facility, www.gbif.org), additional progress is still needed in providing and exploring the utility of different environmental datasets for modeling geographic distributions. In particular, it is unknown if currently available and widely used environmental datasets are sufficient and optimal for modeling distributions of terrestrial species.

The generation and projection of species distribution models requires data layers of environmental information that provide discriminatory power regarding presence and absence of species. As we typically do not know the true distribution of a species, it can be challenging to determine when an appropriate set of environmental variables has been chosen. Ideally, these variables should have direct relevance to ecological or physiological processes determining species distributions, but for many species this information is not generally available (Alvarado-Serrano and Knowles 2014). Correlative niche modeling approaches that rely on statistical associations between species occurrences and environmental variables are frequently used (Peterson et al. 2011, Alvarado-Serrano and Knowles 2014), in which the environmental determinants of habitat suitability are not known *a priori*. The 19 bioclimatic variables from WorldClim (Hijmans et al. 2005) are perhaps the most broadly employed set of environmental data layers for this purpose, on account of their high resolution, global coverage, and availability for both historical and future climate scenarios. However, the biological suitability of these bioclimatic variables and other such environmental datasets for modeling the distribution of the species in question is often not thoroughly assessed.

In the absence of specific knowledge about the environmental variables most likely to determine species distributions, it may be tempting to construct models using a large number of predictor variables, but such models run the risk of poor performance. For example, models built with several highly collinear variables are at an increased risk of overfitting and overparameterization (Dormann et al. 2012, Wright et al. 2014), and may behave unexpectedly when projected to other time periods or geographic regions where they may encounter combinations of variables that have no analog in model training (Dormann et al. 2012, Owens et al. 2013, Warren et al. 2014). Additionally, whether large sets of environmental variables or smaller subsets of environmental data are used can greatly impact model predictions (Rödder et al. 2009, Synes and Osborne 2011, Braunisch et al. 2013). Variable reduction approaches can reduce model overfitting and improve model transferability (Warren et al. 2014, Wright et al. 2014), yet the relative merits of various approaches are poorly characterized and continue to be explored (Araújo and Guisan 2006, Braunisch et al. 2013). In general, variables may be reduced either statistically, or by selecting variables from ecological theory that are likely to be important given the physiology of the organism in question (Kearney et al. 2008, Doswald et al. 2009, Rödder et al. 2009, Synes and Osborne 2011).

Given the recognized importance of variable selection in constructing ecological niche models (Synes and Osborne 2011, Braunisch et al. 2013), increasing the availability of easily accessible datasets of environmental variables that may be ecologically and physiologically important to a variety of organisms should be a priority for improving flexibility and performance of SDM. Several environmental datasets are already available with which to perform SDM (e.g., WorldClim (Hijmans et al. 2005), PRISM (www.prism.oregonstate.edu), ClimateWNA (Wang et al. 2012, Hamann et al. 2013)), but not all of these datasets are transferable among time periods or geographic regions or easily integrated with other variables. Additional environmental data layers that conceptually complement and are formatted for easy use alongside the 19 bioclimatic variables from WorldClim (Hijmans *et al.* 2005) – one of the most widely used environmental datasets for SDM – would broaden the options available for selection of environmental variables (whether based on ecological theory or through statistical variable reduction) and may lead to improved model performance for some species. Despite the description in the literature of formulae for many such variables that could be computed for particular regions or time periods (see Synes and Osborne 2011 as an example), the use of such variables is limited to those researchers with the GIS skills necessary to generate these datasets and the desire to assemble them from several disparate sources.

To help satisfy this need, we introduce the envirem dataset (**ENVI**ronmental **R**asters for **E**cological **M**odeling): specifically, we provide a set of biologically relevant climatic and topographic variables (all of which have previously been described in the literature) at multiple resolutions and time periods. The variables we include were selected in particular because we hypothesize they are likely to have direct relevance to ecological or physiological processes determining distributions of many species. They should therefore facilitate ecologically-informed variable selection, and may also result in improved model performance using statistical variable-thinning approaches. As these variables are intended to complement the existing WorldClim dataset (Hijmans et al. 2005), we provide the envirem dataset at the same extents and resolutions as WorldClim, for the present, mid-Holocene, and Last Glacial Maximum (LGM).

We also provide an R package (R Core Team 2016) that will enable users to generate these variables from primary sources for any resolution, geographic area, or time period, including for future time periods of interest (for which we have not provided static rasters due to the large number of climate change models in existence that are continually updated as climate-change projections improve). Finally, through several case studies, we show that the envirem variables can improve model performance and be valuable additions to the set of variables that are currently widely used in species distribution modeling.

## Methods

We compiled a list of biologically relevant climatic variables (Table 1) that could be derived from monthly temperature and precipitation data (WorldClim v1.4, Hijmans et al. 2005) and monthly extraterrestrial solar radiation (available from www.cgiar-csi.org). These variables are described by Thornthwaite (1948), Daget (1977), Hargreaves et al. (1985), Willmott and Feddema (1992), Vörösmarty et al. (2005), Zomer et al. (2006, 2008), Sayre et al. (2009), Metzger et al. (2013) and Rivas-Martínez and Rivas-Sáenz (2016). We additionally produced two elevation-derived topographic variables, terrain roughness index (Wilson et al. 2007) and topographic wetness index (Boehner et al. 2002, Conrad et al. 2015), generated from a global 30 arc-second elevation and bathymetry digital elevation model (Becker et al. 2009). All variables were produced at the same resolutions as the bioclimatic variables that are currently available through WorldClim: 30 arc-seconds, and 2.5, 5 and 10 arc-minutes. Topographic variables were produced at a 30 arc-second resolution, and subsequently coarsened to match the lower resolutions, rather than constructed directly from lower-resolution elevation data. As such, the topographic variables of large grid cells at coarser scales represent the average fine-scale (i.e., 30 arc-second) values within each grid cell. Calculating the topographic variables in this manner was particularly important to avoid loss of information regarding terrain roughness index when scaling up to coarser resolutions.

**Table 1:** Summary of the variables in the envirem dataset. Citations for variable sources are as follows: A: Zomer et al. (2006, 2008); B: Hargreaves et al. (1985); C: Thornthwaite (1948); D: Willmott and Feddema (1992); E: Vörösmarty et al. (2005); F: Sayre et al. (2009); G: Rivas-Martínez and Rivas-Sáenz (2016); H: Daget (1977); I: Metzger et al. (2013); J: Wilson et al. (2007); K: Boehner et al. (2002); L: Conrad et al. (2015).

| variable abbreviation | brief description | units | source |
| --- | --- | --- | --- |
| annualPET | annual potential evapotranspiration: a measure of the ability of the atmosphere to remove water through evapotranspiration processes, given unlimited moisture | mm / year | A, B |
| aridityIndexThornthwaite | Thornthwaite aridity index: Index of the degree of water deficit below water need | - | C |
| climaticMoistureIndex | a metric of relative wetness and aridity | - | D, E |
| continentality | average temp. of warmest month - average temp. of coldest month | °C | F, G |
| embergerQ | Emberger's pluviothermic quotient: a metric that was designed to differentiate among Mediterranean type climates | - | H |
| growingDegDays0 | sum of mean monthly temperature for months with mean temperature greater than 0°C multiplied by number of days | - | I |
| growingDegDays5 | sum of mean monthly temperature for months with mean temperature greater than 5°C multiplied by number of days | - | I |
| maxTempColdestMonth | max. temp. of the coldest month | °C * 10 | I |
| minTempWarmestMonth | min. temp. of the coldest month | °C * 10 | I |
| monthCountByTemp10 | count of the number of months with mean temp greater than 10°C | months | I |
| PETColdestQuarter | mean monthly PET of coldest quarter | mm / month | I |
| PETDriestQuarter | mean monthly PET of driest quarter | mm / month | I |
| PETseasonality | monthly variability in potential evapotranspiration | mm / month | I |
| PETWarmestQuarter | mean monthly PET of warmest quarter | mm / month | I |
| PETWettestQuarter | mean monthly PET of wettest quarter | mm / month | I |
| thermInd | compensated thermicity index: sum of mean annual temp., min. temp. of coldest month, max. temp. of the coldest month, * 10, with compensations for better comparability across the globe | °C | F, G |
| tri | terrain roughness index | - | J |
| topoWet | SAGA-GIS topographic wetness index | - | K, L |

We generated rasters for all variables at multiple spatial resolutions for current climatic conditions, the mid Holocene (approximately 6,000 years ago) and the Last Glacial Maximum (LGM, approximately 22,000 years ago). For the paleoclimate datasets, we generated variables from three global general circulation models: the Community Climate System Model version 4 (CCSM4, Collins et al. 2006), the Model for Interdisciplinary Research On Climate (MIROC-ESM, Hasumi and Emori 2004), and the model of the Max Planck Institute for Meteorology (MPI-ESM-P, Stevens et al. 2013). As the formulae for some variables require mean monthly temperature, which is available from the WorldClim dataset in the present but not for other time periods, we calculated mean monthly temperature in all time periods as the mean of the maximum and minimum temperatures. In the present, this calculation is highly correlated with the available mean monthly temperatures (Pearson correlation coefficient > 0.99). All raster manipulation and variable creation was carried out in R with the raster package 2.5-2 (Hijmans 2015).

Additional variables derived from and complementing the 19 bioclimatic variables from WorldClim (Hijmans et al. 2005) will only be of value in SDM applications if they represent information not currently contained in the 19 bioclimatic variables. To assess the degree of novelty of these new variables, we calculated the Pearson correlation coefficient between each of the envirem variables and the 19 bioclimatic variables from WorldClim, at a global scale (10 arc-minute resolution), and also by biogeographic realm (Olson et al. 2001, Table 2), for both the present and the past (CCSM global circulation model). Similarly, we also calculated correlation coefficients between terrain roughness index and topographic wetness index with elevation (Table 3) to explore whether these variables contain topographic information not captured by elevation alone.

**Table 2:** Pearson correlation coefficients between each of the climatic envirem variables and the WorldClim bioclimatic variable with which the envirem variable is most strongly correlated (Table S1), globally and in separate biogeographic realms. For each variable and realm, the bottom-left triangle contains the correlation coefficient in the present, and the top-right triangle contains the correlation coefficient in the LGM for the same bioclimatic variable. Grey shading indicates that the absolute value of the correlation is ≤ 0.85.

|  | neotropical | palearctic | nearctic | indo-malay | afrotropic | oceania | australasia | global |
| --- | --- | --- | --- | --- | --- | --- | --- | --- |
| annualPET | 0.88 / 0.93 | 0.94 / 0.93 | 0.96 / 0.94 | 0.8 / 0.85 | 0.9 / 0.93 | 0.83 / 0.55 | 0.94 / 0.93 | 0.91 / 0.94 |
| aridityIndexThornthwaite | -0.84 / -0.81 | -0.78 / -0.62 | -0.73 / -0.25 | 0.89 / 0.85 | -0.83 / -0.79 | -0.8 / -0.71 | -0.91 / -0.86 | -0.81 / -0.67 |
| climaticMoistureIndex | 0.93 / 0.91 | -0.82 / -0.83 | -0.59 / -0.63 | 0.91 / 0.91 | 0.98 / 0.98 | 0.89 / 0.88 | 0.95 / 0.96 | 0.81 / 0.6 |
| continentality | 1 / 1 | 1 / 1 | 0.99 / 0.99 | 0.99 / 1 | 0.99 / 0.99 | 1 / 1 | 1 / 1 | 1 / 1 |
| embergerQ | 0.95 / 0.95 | 0.91 / 0.9 | 0.94 / 0.87 | 0.88 / 0.85 | 0.94 / 0.95 | 0.92 / 0.85 | 0.97 / 0.94 | 0.93 / 0.91 |
| growingDegDays0 | 1 / 1 | 0.93 / 0.87 | 0.89 / 0.77 | 1 / 1 | 1 / 1 | 1 / 1 | 1 / 1 | 0.97 / 0.94 |
| growingDegDays5 | 1 / 0.99 | 0.91 / 0.84 | 0.85 / 0.74 | 1 / 0.99 | 1 / 1 | 1 / 1 | 1 / 0.99 | 0.96 / 0.92 |
| maxTempColdest | 0.98 / 0.98 | 1 / 1 | 1 / 0.99 | 0.97 / 0.97 | 0.92 / 0.93 | 0.98 / 0.97 | 0.97 / 0.97 | 1 / 1 |
| minTempWarmest | 0.98 / 0.98 | 0.99 / 0.99 | 0.98 / 0.99 | 0.96 / 0.93 | 0.96 / 0.95 | 1 / 0.98 | 0.96 / 0.97 | 0.98 / 0.99 |
| monthCountByTemp10 | 0.87 / 0.88 | 0.95 / 0.89 | 0.93 / 0.81 | 0.74 / 0.86 | 0.49 / 0.73 | 0.62 / 0.77 | 0.7 / 0.92 | 0.95 / 0.94 |
| PETColdestQuarter | 0.93 / 0.95 | 0.87 / 0.87 | 0.87 / 0.77 | 0.91 / 0.94 | 0.82 / 0.85 | 0.58 / 0.52 | 0.93 / 0.94 | 0.9 / 0.89 |
| PETDriestQuarter | 0.83 / 0.84 | 0.92 / 0.91 | 0.87 / 0.87 | 0.89 / 0.9 | 0.84 / 0.87 | -0.74 / -0.57 | 0.74 / 0.73 | 0.87 / 0.87 |
| PETseasonality | 0.98 / 0.97 | 0.73 / 0.85 | 0.91 / 0.97 | 0.97 / 0.94 | 0.93 / 0.94 | -0.76 / -0.58 | 0.96 / 0.95 | 0.7 / 0.48 |
| PETWarmestQuarter | 0.74 / 0.8 | 0.98 / 0.98 | 0.99 / 0.98 | 0.94 / 0.94 | 0.84 / 0.88 | 0.84 / 0.7 | 0.94 / 0.9 | 0.91 / 0.95 |
| PETWettestQuarter | 0.79 / 0.85 | 0.89 / 0.92 | 0.95 / 0.94 | 0.68 / 0.76 | 0.62 / 0.76 | -0.82 / -0.1 | 0.91 / 0.93 | 0.83 / 0.91 |
| thermicityIndex | 1 / 0.99 | 0.98 / 0.98 | 0.98 / 0.99 | 0.96 / 0.97 | 0.99 / 0.99 | 1 / 1 | 0.97 / 0.98 | 0.98 / 0.99 |

**Table 3:** Pearson correlation coefficients between envirem topographic variables and elevation, at a global scale as well as in different biogeographic realms.

|  | neotropical | palearctic | nearctic | indo-malay | afrotropic | oceania | australasia | global |
| --- | --- | --- | --- | --- | --- | --- | --- | --- |
| terrain roughness | 0.65 | 0.58 | 0.48 | 0.83 | 0.41 | 0.19 | 0.65 | 0.46 |
| topographic wetness | -0.59 | -0.45 | -0.42 | -0.67 | -0.37 | -0.49 | -0.53 | -0.39 |

## Case Studies

To investigate how the inclusion of the envirem variables could affect the performance and predictions of species distribution models, we generated species distribution models with Maxent v3.3.3k (Phillips et al. 2006) for 20 North American terrestrial vertebrate species, using the curated occurrence dataset from Waltari et al. (2007). Specifically, we generated niche models using three different sets of initial environmental predictor variables. Firstly, we generated models using only the 19 bioclimatic variables from WorldClim (referred to hereafter as the *bioclim* model). Secondly, we built models using the 19 bioclimatic variables plus 14 of the climatic envirem variables (hereafter referred to as the *bioclim + envirem-clim* model).

Finally, we generated niche models with the 19 bioclimatic variables and 16 envirem variables, including 14 climatic variables and the two topographic variables (the *bioclim + envirem-all* model). Note that none of the models, including *bioclim + envirem-all*, included elevation as a predictor variable. We chose not to include two variables, aridityIndexThornthwaite as it was conceptually redundant with the climaticMoistureIndex, and monthCountByTemp10 because it is a categorical variable that would not have been amenable to the variable selection procedure that we applied. Finally, we did not generate any models using only the envirem variables without the 19 bioclimatic WorldClim variables, as the envirem variables are intended to supplement, not replace, the bioclimatic variables. All distribution modeling was performed in the dismo package v1.0-15 in R (Hijmans et al. 2016) from rasters at a 2.5 arc-minute resolution.

To construct each model, we first spatially thinned the occurrence records, retaining only occurrences that were greater than ten kilometers in proximity to one another, using the spThin package in R (Aiello-Lammens et al. 2015). For each species individually, we defined the model-training region by adding a 1,000 km buffer around all occurrence records (Supplementary Figure S1). All occurrence data and rasters were transformed and projected to the North America Albers Equal Area Conic projection, as it has been shown that a failure to account for changing grid-cell area across latitudes can negatively impact SDM results (Budic et al. 2015). We statistically thinned variables to include in each model for each species using the “corSelect” function in the fuzzySim package v1.6.3 in R (Barbosa 2015) where each pair of variables that is correlated above a set threshold is tested against the response variable (species presence and absence) with a bivariate model. The variable with a better fit as measured with AIC is selected while the other is dropped, and the procedure is repeated until all pairwise correlations are below the threshold. We applied a correlation threshold of 0.75, and generated pseudo-absences from 10,000 randomly sampled points throughout the training region (excluding grid cells with known occurrence records) because there were no true absence records in our data.

For each species, we measured SDM performance for the *bioclim*, the *bioclim + envirem-clim* and the *bioclim + envirem-all* models (with reduced sets of variables via statistical thinning as described above, Table 4) using three threshold-independent evaluation metrics: AUC_TEST,_ AUC_DIFF_, and the size-corrected Akaike Information Criterion (AICc). AUC_TEST_ is a metric that measures the discriminatory ability of the species distribution model at test localities withheld during model construction, and thus represents the ability of the model to predict species presence (Peterson et al. 2011). AUC_DIFF_ is the difference between the AUC calculated from training localities and AUC_TEST_, and is a measure of model overfitting, with higher values of AUC_DIFF_ representing more overfit models (Warren and Seifert 2011). AICc is an information theoretic metric that balances model fit against degrees of freedom from parameterization (i.e., model complexity), such that lower values of AICc correspond to models with better goodness-of-fit accounting for model complexity (Burnham and Anderson 2004, Warren and Seifert 2011). For AUC metrics, we partitioned calibration and evaluation data via the masked geographically-structured partitioning scheme described by Radosavljevic and Anderson (2014), implemented in the R package ENMeval v0.2.1 (Muscarella et al. 2014), which leads to more realistic and less biased estimates of SDM performance than the more traditionally used random *k*-fold partitioning scheme. This partitioning scheme divides occurrence records into four geographic regions with an equal number of occurrence records, and calculates AUC metrics as the average of those metrics calculated individually using each of the four possible partitions of geographic regions into one region of evaluation data and three regions of calibration data. AICc was calculated from the full, non-partitioned models.

**Table 4:**
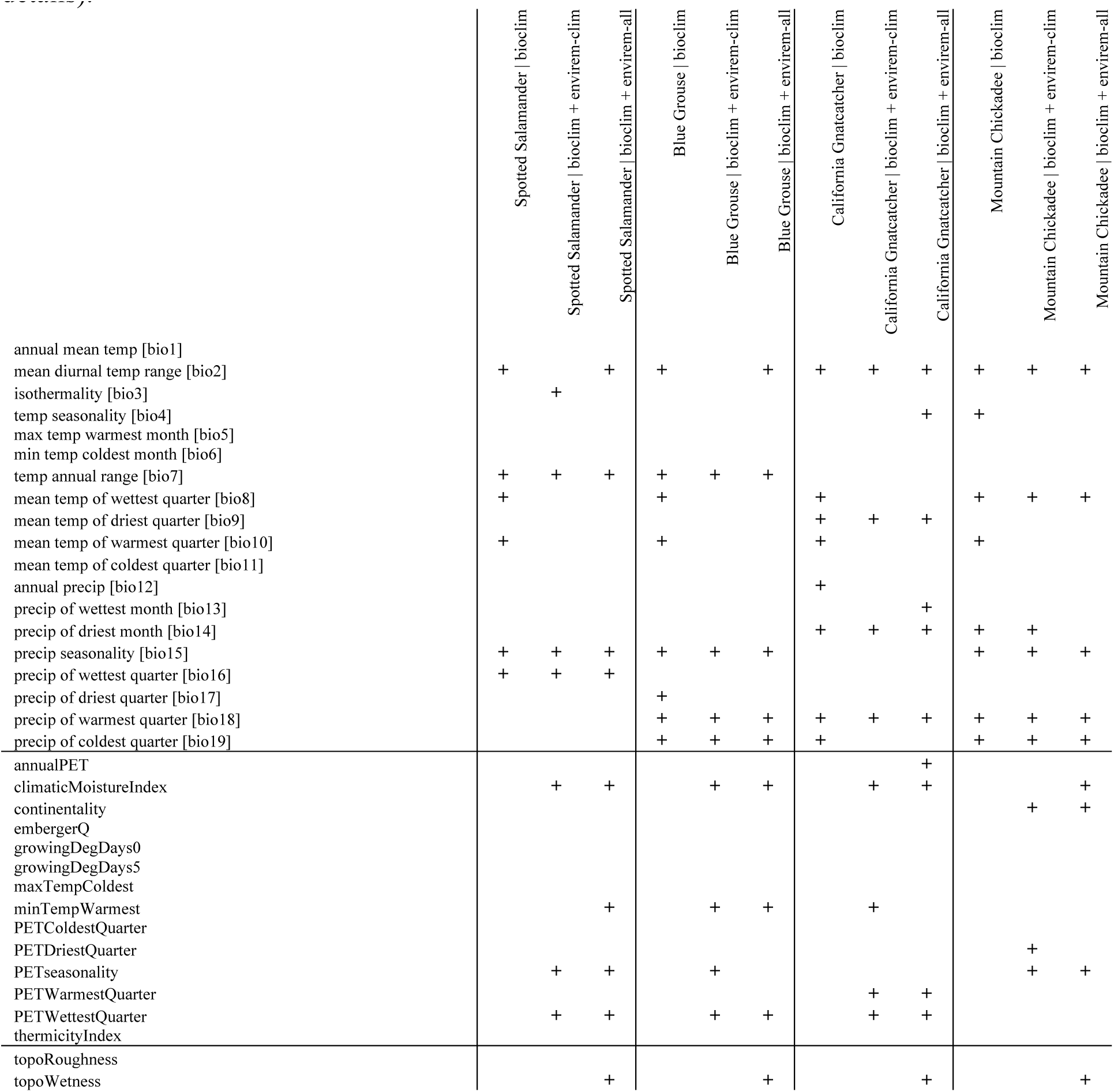
envirem and WorldClim variables included in the *bioclim*, *bioclim + envirem-clim* and *bioclim + envirem-all* models, for four case study species. Variables included in each model were selected using a statistical variable selection approach (see Methods section for additional details).

The complexity of SDMs built with Maxent can be adjusted with the regularization multiplier, increased values of which lead to less parameterized models, as well as with the inclusion of additional feature classes (i.e., transformations of the original predictor variables) that allow for increasingly complex models. We evaluated distribution models across different sets of permissible feature classes, and for each of these, across a range of regularization multiplier values. The evaluation metrics described above were used to determine optimal feature class and model complexity for each model individually (Muscarella et al. 2014).

After selecting optimal feature class and model complexity for each model, we also compared performance of the optimal models across each of the three variable sets (i.e., *bioclim*, *bioclim + envirem-clim*, and *bioclim + envirem-all*) using the same evaluation metrics. The AUC metrics describe absolute performance of the models (ranging from 0 to 1). AICc, however, describes relative performance of candidate models. For this metric, we define a model as having substantial support over another if it has a difference in AICc greater than or equal to four, as models with AICc values more similar than this are generally considered to have equivalent support (Burnham and Anderson 2004). Although we present results for all evaluation metrics, we ultimately favor AICc for selecting the optimal model and variable set for each species, as the focus of our case studies is on model comparison, and AICc has been shown to perform better than AUC metrics according to a range of criteria, including the selection of optimal levels of model complexity, model transferability in space and time, and the relative ranking of variable importance (Warren and Seifert 2011, Warren et al. 2014, Moreno-Amat et al. 2015).

The impact of using different environmental variables in niche modeling may not be apparent if two sets of variables lead to similar projected distributions in the present. However, if the degree of correlation between two different sets of variables differs in the past compared to in the present, then variable choice might have a greater effect on SDM projections to other time periods. To explore this possibility, we calculated niche similarity in the present and in the LGM using Schoener’s *D* (Schoener 1968, Warren et al. 2008), a metric that quantifies the degree of niche overlap in geographic space. Values of *D* range from 0 (completely different niches across geographic space) to 1 (identical niches over geographic space). Overlap was quantified with the fuzzySim package in R (Barbosa 2015). For each case-study species we focused the niche overlap calculation on the geographic regions of the model projections where comparisons among models are most meaningful, rather than across broad regions of the continent where all models predict low habitat suitability and are thus very similar. In particular, we calculated niche overlap statistics only over the geographic region predicted to contain suitable habitat in at least one of the models. To define this region, we first reduced the geographic extents of interest for both the projected *bioclim* and *bioclim + envirem-clim* models individually using a habitat suitability threshold that preserved 95% of the training presences. We further excluded areas outside the model training region, except for a few species where the majority of the predicted LGM distribution lay outside the training region. Finally, we combined these regions for both the *bioclim* and *bioclim + envirem-clim* models and calculated niche overlap from (non-thresholded) model projections within this combined region.

We did not project the *bioclim + envirem-all* model to the LGM, because topographic variables are difficult to interpret for the LGM in glaciated regions of North America. These regions have experienced substantial changes in topography since the LGM due to glacial erosion (Bell and Laine 1985). However, we note that models using topographic variables could be projected to the LGM in particular regions of interest where topographic variables can be assumed to have remained static since the LGM (e.g., unglaciated regions of California, Bemmels et al. 2016).

## Results

The envirem dataset comprises variables that were generated for three time periods (present, mid-Holocene and the LGM), using several different general circulation models (CCSM4, MIROC-ESM, MPI-ESM-P) at multiple resolutions, so as to facilitate integration with rasters from WorldClim (Hijmans et al. 2005). All rasters are available for download at envirem.github.io. To enable users to generate these variables from other circulation models or time periods, we have provided all code in an R package “envirem”, available from CRAN.

At a global scale, most new climatic variables were highly correlated with at least one of the 19 bioclimatic variables from WorldClim (Table 2). The aridity-related variables (i.e., climatic moisture index and Thornthwaite’s aridity index) and some of the PET-related variables were the least redundant at the global scale. However, many of the new variables were less highly correlated with the 19 bioclimatic variables within specific biogeographic realms. Oceania and the Afrotropics were the realms with the greatest number of new variables with lower maximum correlation coefficients (≤ 0.85), indicating that niche models of species from those regions may benefit most from the inclusion of these new variables. More often than not, correlations were lower during the LGM than the present, which indicates that even if specific sets of variables are redundant in the present, they may not necessarily be redundant in other time periods and variable choice could have greater impacts on model projections to other time periods. All new climatic variables had a maximum correlation of ≤ 0.85 in at least one biogeographic realm during at least one time period, with the exception of continentality, thermicity index, maximum temperature of the coldest month and minimum temperature of the warmest month. Some new variables were consistently most highly correlated with the same bioclimatic variable from WorldClim across regions, while other new variables were most highly correlated with different bioclimatic variables across different regions (Table S1).

In terms of topographic variables derived from elevation, terrain roughness index was not highly correlated with elevation globally or in any biogeographic region (Table 3). Topographic wetness index was also not highly correlated with elevation (Table 3), even though higher values of topographic wetness are conceptually associated with lower elevations at a local scale (i.e., within a given watershed; Boehner et al. 2002).

## Case studies

Statistical thinning of the sets of variables prior to ecological niche modeling substantially reduced the number of variables, with three to 11 variables retained in each model (Table 4, Supplementary Table S2). For all species, at least one envirem variable was retained in the *bioclim + envirem-clim* models. For the *bioclim + envirem-all* models, at least one topographic variable was retained for 19 of 20 species. For most species, one or more bioclimatic variables that were retained in the *bioclim* model were dropped from the *bioclim + envirem-clim* and *bioclim + envirem-all* models and were replaced by one or more of the envirem variables, indicating that these variables are more strongly predictive of the presence and absence of the species than the dropped bioclim variables (Table S2). The impact of including envirem variables on model performance varied among species, but models containing envirem variables performed substantially better (according to the AICc metric) than the *bioclim* model in 17 of 20 species.

In Figure 1, we highlight results for four species that show particularly distinct improvement with the envirem variables: the spotted salamander (*Ambystoma maculatum*), the Blue Grouse (*Dendragapus obscurus*), the California Gnatcatcher (*Polioptila californica*) and the Mountain Chickadee (*Poecile gambeli*). In these four species, inclusion of envirem variables led to improvements in all metrics of model performance, although differences in AICc values were more substantial than differences in AUC metrics for these species. Across the 16 other case study species (Figure S2), an improvement in performance when including envirem variables was found for ten species according to greater AUC_TEST_ values (*Dicamptodon tenebrosus*, *Dicrostonyx groenlandicus*, *Eumeces fasciatus*, *Glaucomys sabrinus*, *Glaucomys volans*, *Lampropeltis zonata*, *Lepus arcticus*, *Martes americana*, *Myodes gapperi* and *Plethodon idahoensis*). However, substantial improvements in model performance (improvement by more than four AICc units) were found for all but three species according to AICc values, with no substantial difference for *Dicamptodon tenebrosus* and a substantial decrease in performance for only two species (*Glaucomys volans and Martes americana*). Inclusion of envirem topographic variables specifically led to especially notable improvements in AICc scores for *Poecile gambeli* (Figure 1), *Dicrostonyx groenlandicus*, *Lepus arcticus*, *Myodes gapperi* and *Plethodon idahoensis* (Supplementary Figure S2).

**Figure 1.**
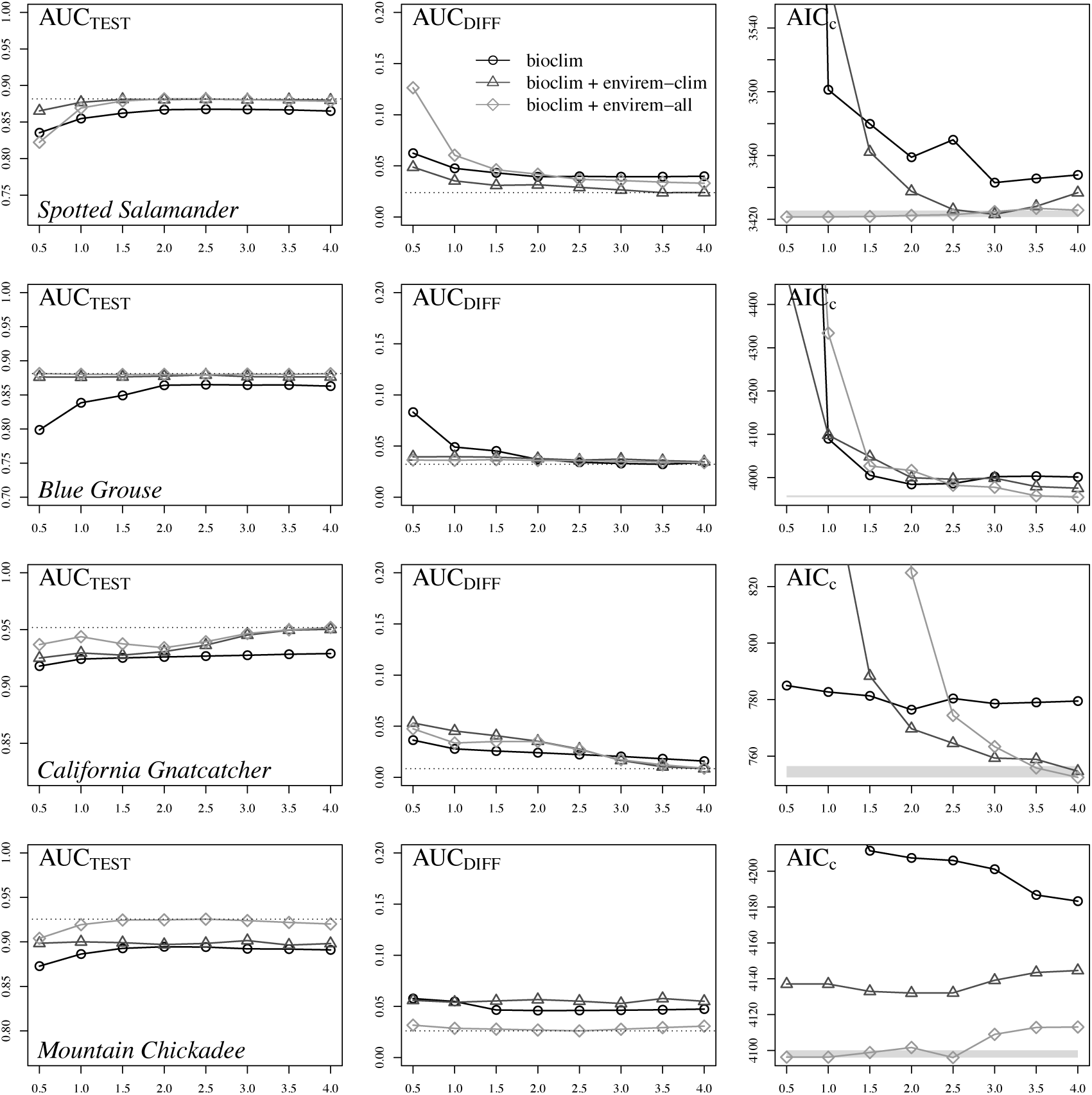
Ecological niche model performance with and without the envirem variables for four selected case study species. Each line represents the set of feature classes that led to the best performance according to either AUC_TEST_ (left and middle panels) or AICc (right panel), with performance evaluated across a range of regularization multiplier values (Supplementary Table S3). In the AUC plots, the dotted line represents the value for the best-performing model. In the AICc plots, the grey shading represents a AICc of 4 from the best (lowest) AICc score. Performance of models within the grey polygon is not considered to be substantially different (Burnham and Anderson 2004).

The optimal Maxent parameters identified by the model evaluation metrics were typically not concordant across the *bioclim*, *bioclim + envirem-clim*, and *bioclim + envirem-all* models (Supplementary Table S3). Similarly, as the different metrics evaluate the niche models using conceptually different criteria, AUC-based evaluations did not identify the same Maxent parameters as AICc-based evaluations (Supplementary Table S3). As the focus of our case studies is on the choice of variables employed, an in-depth examination of the differences between AUC and AICc-based optimization of Maxent is beyond the scope of our study. We therefore focus the rest of our results and discussion on comparing predictions of models that were optimized based on AICc (see Methods).

Projections of the AICc-optimized species distribution models constructed with and without the envirem variables generally did not differ greatly at continental scales for the current time period, but regional-scale differences in habitat suitability were observed. For the four case-study species showing greatest improvement in all evaluation metrics, the overall suitable ranges are very similar, though not identical, at the continental scale (Figure 2). In finer-scale maps focusing on a particular region of interest, however, there are more substantial differences in suitability across the landscape at a regional scale (Figure 2). For example, suitability of the California Central Valley for *Polioptila californica* is much higher in the *bioclim* model than in the *bioclim + envirem-clim* model. Similarly, regions of the California coast and northwestern Great Basin for *Dendragapus obscurus* are also considerably different across models, as well as large areas of the interior range of *Poecile gambeli*. Niche overlap (Schoener’s *D*) between the two models averaged 0.88 for these four species and 0.9 across all modelled species (Figure S3, Table S4).

**Figure 2.**
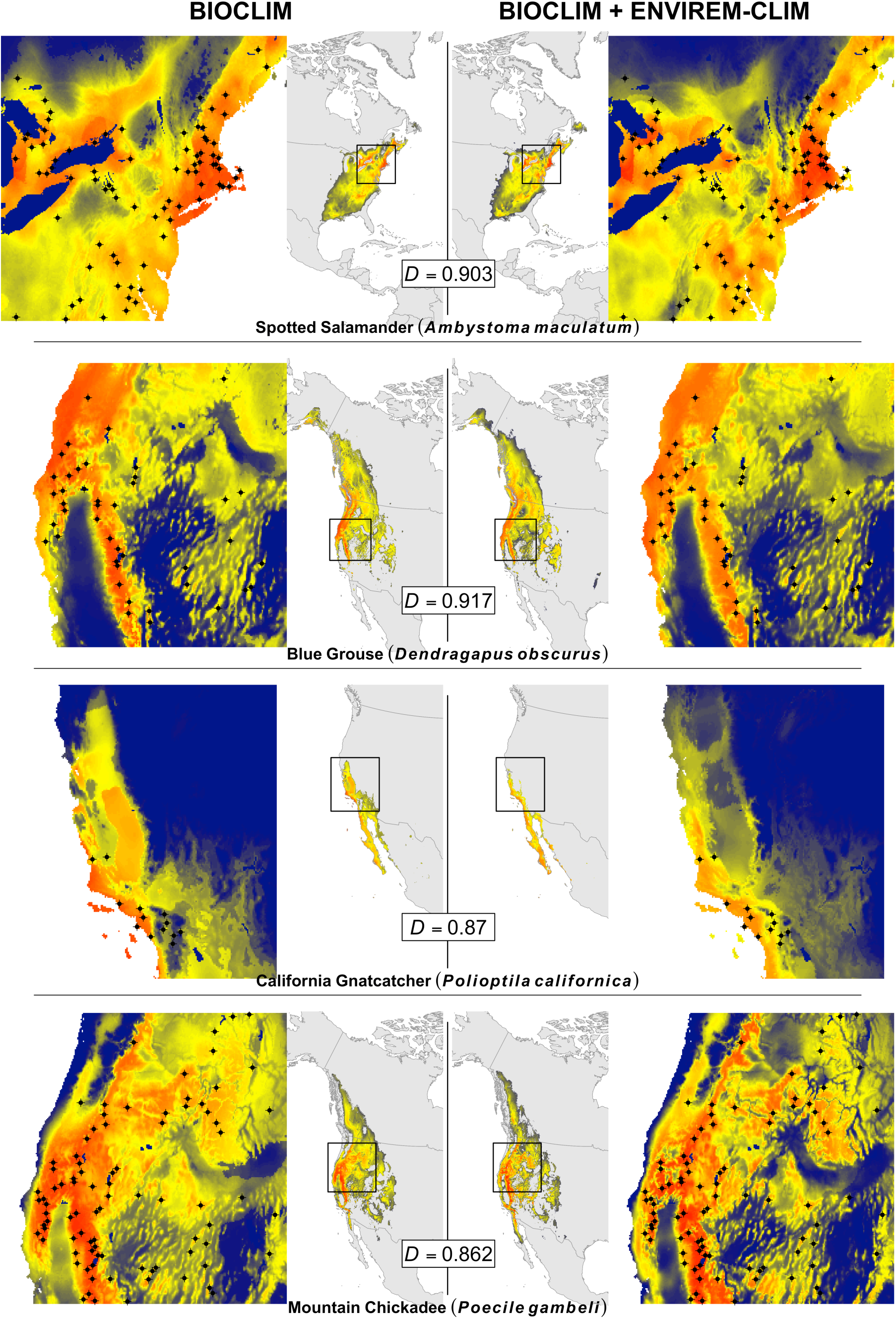
Predicted habitat suitability during the current time period for four case study species, from Maxent models optimized in terms of feature class and regularization parameter according to the AICc metric, for models constructed with and without the envirem variables. Suitability scores range from 0 (blue) to 1 (red). The central, continental-scale maps show habitat suitability within the training region only (see text for explanation), with predicted habitat suitability below a 95% training presence threshold considered to be unsuitable (grey). The outer maps show detail from the region within the box on the continental maps, selected to highlight local-scale differences between the models. Occurrence records are shown as black points. Schoener’s *D* niche overlap is calculated between the *bioclim* and the *bioclim + envirem-clim* models, exclusively within the thresholded training regions (Supplementary Figure S1; see the Methods section for additional details).

Differences between the predictions of the AICc-optimized *bioclim* and *bioclim + envirem-clim* models become more pronounced when projected to the LGM (Figure 3, Supplementary Table S4). In particular, Schoener’s *D* niche overlap scores are much lower in the LGM (mean = 0.72) compared to the present, and for many species there are considerable differences between models in predicted distribution in the LGM (Figure 3). For *Ambystoma maculatum*, habitat suitability in the *bioclim* model was highest on exposed continental shelf off the coast of North Carolina, whereas in the *bioclim + envirem-clim* model the highest habitat suitability was in the Lower Mississippi River Valley. For *Dendragapus obscurus*, connectivity between regions was greater in the *bioclim + envirem-clim* model, and areas of high habitat suitability included the Columbia Plateau and northern Cascades. Both models for this species also showed marginally to moderately suitable habitat in western Canada and Alaska, although this may be an overprediction as at least part of this region was covered by the Cordilleran ice sheet during the LGM (Dyke et al. 2002). For *Polioptila californica*, the *bioclim* model predicted large regions of California to be suitable, including California’s Central Valley, whereas in the *bioclim* + *envirem-clim* model, higher suitability was primarily restricted to Baja California and coastal regions of southern California. For *Poecile gambeli*, visual differences between model projections were even greater, with high habitat suitability in the Rocky Mountains in the *bioclim + envirem-clim* model only, and much higher habitat suitability throughout most of the species’ range overall, and the Great Basin in particular.

**Figure 3.**
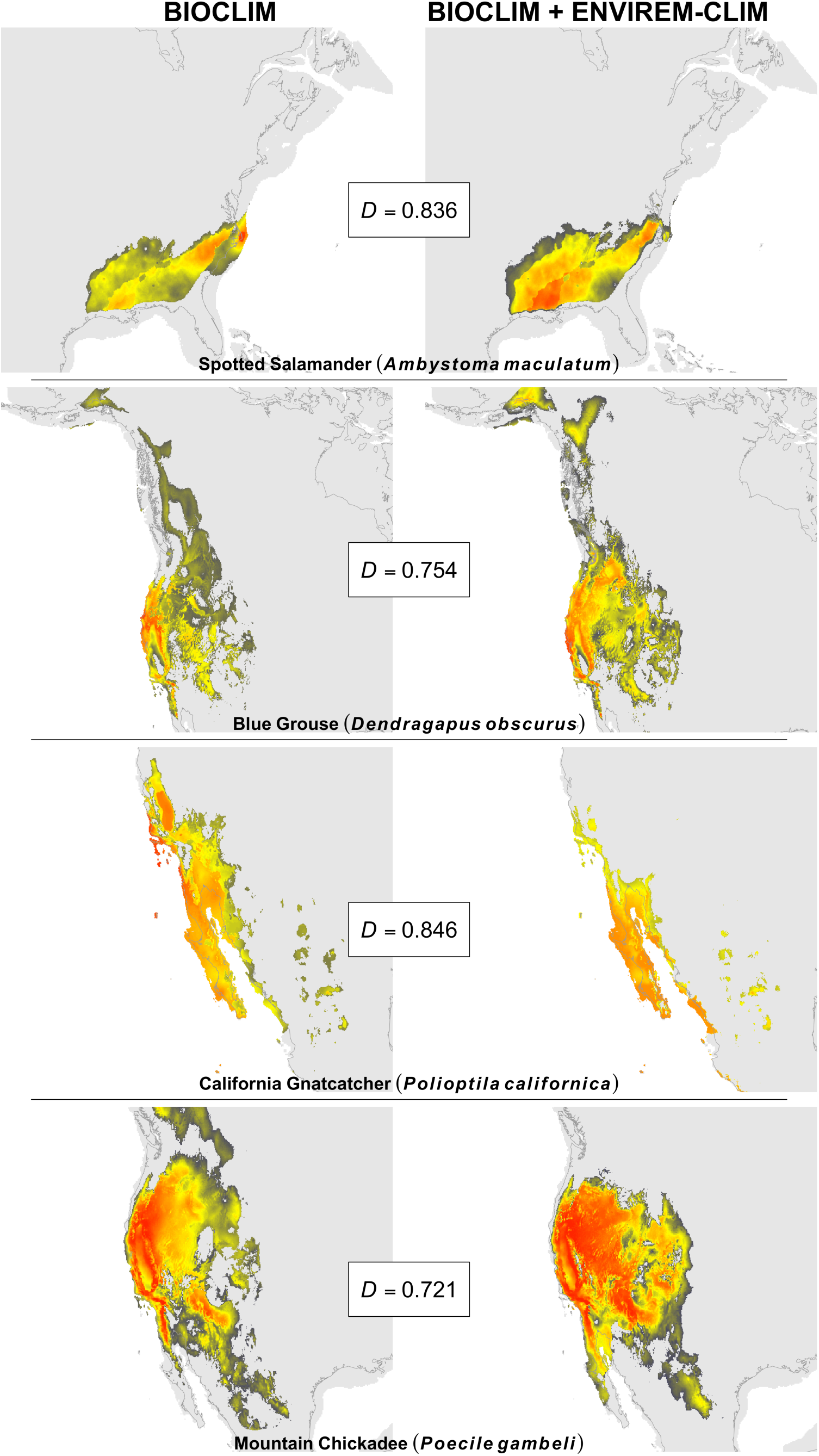
Predicted habitat suitability during the Last Glacial Maximum for four case study species, for models constructed with and without the envirem variables. Suitability scores range from 0 (blue) to 1 (red). Optimization of model parameters and thresholding are as in Figure 2. Schoener’s *D* niche overlap is calculated between the *bioclim* and the *bioclim + envirem-clim* models, exclusively within the thresholded training regions (Supplementary Figure S1; see the Methods section for additional details). Habitat suitability is shown within the training region only, with predicted habitat suitability below a 95% training presence threshold considered to be unsuitable (grey).

## Discussion

We have generated 18 climatic and topographic variables that will be valuable in a broad array of applications for species distribution modelling, and have made these variables easily available and complementary to an existing widely-used environmental dataset. Although they are largely derived from the same underlying dataset as the bioclimatic variables from WorldClim, we have demonstrated that including the envirem variables in SDM can lead to notable improvements in performance and differences in projections of species distribution models. Inclusion of these new variables led to substantial improvement in SDM performance (AICc metric) in 17 out of 20 species, and substantially worse performance in only two species. Although inclusion of the envirem variables did not always lead to significantly improved performance, the fact that they were beneficial to many species indicates that they are generally worth consideration when constructing species distribution models. The species-specific nature of our results also highlight the importance of following best practices for variable selection and parameter optimization, as we have done here. The importance of particular variables in SDM will be a function of the species under study, its distribution in geographic and climatic space, the time period and geographic region of interest, and the ultimate question being addressed.

Nonetheless, the links to ecological and physiological processes represented in many of the ENVIREM variables mean that they will likely be particularly useful for a wide variety of applications.

### Potential Applications

As we have showcased here, the envirem dataset will be of immediate value in SDM applications and will potentially lead to the generation of better species distribution models. If variable selection is done via statistical approaches, then inclusion of these variables will allow researchers to start with a larger pool of biologically relevant options, thereby increasing the odds that variables that are highly informative regarding the presence and absence of a species will be discovered. If the goal is to select variables *a priori* based on the ecology and natural history of the organism, then the envirem variables will provide valuable options, as they are likely to be ecologically relevant to certain species and may have specific ties to biological processes for many species (see below). SDM has been employed as a tool in a large variety of studies, and the inclusion of new variables has the potential to impact their conclusions.

Identifying better sets of predictor variables for certain species could, among other things, potentially alter projections of species’ invasiveness for particular regions (Peterson and Nakazawa 2008), alter our understanding of potentially suitable habitat for species introductions (Martínez-Meyer et al. 2006), lead to identification of new areas of high habitat suitability for conservation interest, affect predictions of shifts in habitat suitability in response to future climate change (Thuiller 2004, Hijmans and Graham 2006, Morin and Thuiller 2009), lead to new phylogeographic hypotheses about where species may have been distributed in the past (Chan et al. 2011, He et al. 2013, Bemmels et al. 2016), and impact our understanding of the evolution of climatic tolerances across related species (Title and Burns 2015, Kozak and Wiens 2016).

With these additional variables, ecologists and evolutionary biologists will also be able to craft more specific hypotheses that are informed by the ecology of the organisms under study.

For example, in an integrative distributional, demographic and coalescent (iDDC) framework (Knowles and Alvarado-Serrano 2010, Brown and Knowles 2012, He et al. 2013), these variables will allow for the specification of competing hypotheses pertaining to the relative importance of different climatic and topographic variables in constraining the distribution of species over time (e.g., Bemmels et al. 2016), giving researchers greater flexibility than currently exists in modeling spatial and genetic patterns over time.

To our knowledge, this is the only existing multi-variable dataset that is truly complementary to WorldClim in its breadth, application and accessibility. The Climond dataset (Kriticos et al. 2011) provides an extended suite of bioclimatic variables only at 10 and 30 arc-minutes for current and future climate scenarios, while the Ecoclimate dataset (Lima-Ribeiro et al. 2015) provides only the standard 19 bioclimatic variables for multiple past, present and future time periods at 30 arc-minutes. Other variables potentially useful for biodiversity modeling have been released, such as habitat heterogeneity (Tuanmu and Jetz 2015), global cloud cover (Wilson and Jetz 2016) and region-specific variables (Wang et al. 2012, Hamann et al. 2013), but these variables are either not transferrable to other time periods, not available globally or not available at finer spatial resolutions. In contrast, the envirem dataset includes additional variables (some of which overlap with the Climond dataset) at all of the resolutions currently available from WorldClim, for past and current time periods. The envirem R package makes it possible to generate these variables for other time periods as well, or from alternative input datasets, allowing users to easily customize their use of these variables.

### Biological relevance of envirem variables

Although the potential applications of these variables to SDM are vast, one unique benefit of the envirem variables is their potential for improving our ability to construct niche models informed by ecological knowledge and natural history. Biologically informed niche models may be constructed for species for which the conceptual relationships between particular variables and biological processes relevant to determining a species’ distribution are known *a priori* (Kearney et al. 2008, Doswald et al. 2009, Rödder et al. 2009, Synes and Osborne 2011), or may be constructed with the intention of exploring and testing different hypotheses about these relationships (e.g., Bemmels et al. 2016).

The potential mechanisms by which the envirem variables may determine distributions are numerous and will be specific to the species of interest. In general, subsets of the envirem variables may directly control species distributions, or (more commonly) may impact other processes that in turn determine distributions (Austin 2002). The particular variables included in the envirem dataset were selected because of their clear conceptual links to particular ecological processes and indices. For example, growing degree-days are predictive of plant phenology and growth rate (e.g., McMaster and Wilhelm 1997), processes which impact species range limits (e.g., Morin et al. 2007) and drive local adaptation (e.g., Howe et al. 2003). Evapotranspiration not only describes climate generally, but is also physiologically linked to plant growth potential due to its impact on gas exchange with the atmosphere and temperature regulation (Thornthwaite 1948, Katul et al. 2012). The more complex climatic indices included in the envirem variables (e.g., thermicity, aridity, moisture, Emberger’s pluviothermic quotient) may characterize environmental conditions that are more directly physiologically relevant to given species than simple descriptors of climate such as temperature or precipitation alone (e.g., Daget 1977).

Finally, the topographic envirem variables could conceivably be important predictors of habitat types associated with local- to regional-scale relief that may be key predictors of species distributions at these spatial scales (e.g., Lassueur et al. 2006, Austin and Van Niel 2011). We have provided just a few examples of potential links to biological factors that could determine species distributions, but the ecological relevance of any of the envirem variables is likely to be species-specific and different species’ distributions may be associated with environmental variables because of different mechanisms. Nonetheless, it is this type of conceptual relevance and these potential links to physiological and ecological processes that will make the envirem variables particularly useful for many SDM applications.

### Incorporating envirem variables into SDM best practices

Ideally, the choice of variables for niche modeling should be informed by knowledge of the natural history and ecology of the organism under study, as this approach has been shown to produce more realistic niche models (Rödder et al. 2009, Saupe et al. 2012). However, it is most often the case that such information is not readily known (Alvarado-Serrano and Knowles 2014). How one should go about choosing bioclimatic variables is still an open question, the impact of which can be considerable (Peterson and Nakazawa 2008, Synes and Osborne 2011, Braunisch et al. 2013). It is generally not considered best practice to include all bioclimatic variables, as they exhibit a high degree of collinearity. This collinearity tends to lead to overly complex, overfit models (Rodda et al. 2011). Additionally, the nature of the correlation between bioclimatic variables may differ across time periods, potentially leading to unexpected behavior in SDM projections (Synes and Osborne 2011, Rodda et al. 2011, Dormann et al. 2012, Warren et al. 2014). While we expect that many researchers will find the envirem variables extremely useful for a variety of applications, we recommend that the merits of including all or some of the ENVIREM variables should be carefully considered relative to the specific application, and that variable thinning, model optimization, and other best practices in ecological niche modeling should be followed (e.g., Merow et al. 2013, Alvarado-Serrano and Knowles 2014). For example, as we do not have in-depth ecological information about the species whose ecological niches were modeled in our case studies, we employed a statistical approach to variable thinning in order to reduce the number of correlated variables, while retaining the variables with the greatest explanatory power.

An important finding of our case studies was that the difference between the *bioclim* and *bioclim + envirem-clim* models, as measured with Schoener’s *D*, was small in the present, but greater in the LGM. Choice of predictor variables has previously been shown to have large impacts on model projections to other time periods or geographic regions (Peterson and Nakazawa 2008, Synes and Osborne 2011, Braunisch et al. 2013). The impact of variable selection points both to the utility of additional variables for developing and testing hypotheses about shifts in species distributions across different time periods and in novel spatial contexts, but also to the need for caution when making modeling decisions. Ideally, models could be evaluated in past time periods with independent fossil occurrences (Davis et al. 2014, Gavin et al. 2014, Moreno-Amat et al. 2015), but their availability will depend on the taxon under study.

In addition to the question of which environmental variables to use, a growing number of studies have demonstrated that species-specific tuning of virtually all steps in the niche modeling pipeline can lead to improved results, and that Maxent’s default behavior is often not sufficient to achieve optimal performance (Anderson and Gonzalez Jr 2011, Warren and Seifert 2011, Merow et al. 2013, Radosavljevic and Anderson 2014, Moreno-Amat et al. 2015). Although we could have held all aspects save the predictor variables constant in the generation of niche models in order to be able to compare the results directly, generating models in this way is considered poor practice. Instead, we chose to independently generate the best possible models, given current best practices. We found that Maxent’s default parameters were rarely optimal (Table S3), which echoes the findings of others that parameter tuning is an important step toward generating less overfit and more transferable species distribution models (Anderson and Gonzalez 2011, Warren and Seifert 2011, Merow et al. 2013, Radosavljevic and Anderson 2014, Moreno-Amat et al. 2015). Different evaluation metrics most often did not lead to the selection of the same optimized parameters (Table S3). This is expected, as AICc is intended to minimize the number of necessary parameters, while AUC metrics are not. Regardless of the environmental variables selected for SDM, the optimization of model parameters should always be considered, as model parameters can have a large impact on model performance and predictions (Figure 2, Figure S2).

### Utility of topographic variables in SDM

In addition to climatic variables, we also generated two topographic indices: topographic roughness and topographic wetness. These variables offer novel information as they are not redundant with elevation (Table 3), an environmental variable which is already broadly available for SDM. The use of elevation in SDM has been controversial (Hof et al. 2012), and may be particularly problematic when projecting to other time periods or geographic contexts where relationships between elevation and the climatic factors determining a species’ niche may be different than the relationships in the context in which the model was built. However, the topographic roughness and topographic wetness indices are less likely to suffer from this complication because they are less causally linked than elevation to regional-scale climate, and they contain topographic information that may be useful for determining species distributions independent of climate. In particular, topographic roughness index may be a reasonable surrogate for habitat heterogeneity and microsite availability that could be relevant to determining geographic distributions of some species, and topographic wetness index may help distinguish between areas that experience similar regional climate but differ markedly in microhabitat due to relative drainage position within a watershed.

However, it is important to consider whether topographic variables are available at an appropriate geographic scale for predicting species distributions. Variation in topographic features associated with microhabitats may occur at a much finer scale than that at which topographic variables are assessed, which could reduce their utility for SDM (Lassueur et al. 2006, Austin and Van Niel 2011, Pradervand et al. 2014). Since all topographic envirem variables at all resolutions are ultimately averaged from values calculated from the finest-scale (30 arc-second) elevational model (see *Methods*), we have minimized concerns about the potential mismatch between the scale at which the indices were generated and at which topography is relevant to a species. However, it is still important to consider whether variation in topographic roughness and wetness at the 30 arc-second scale (approximately 926 m at the equator) is likely to be meaningful for the species in question for the particular geographic region of interest and intended modeling application.

Nonetheless, our case studies revealed that including topographic variables led to distinct improvement in SDM performance for several species, in some cases significantly exceeding the improvement gained by adding only the climatic envirem variables (Figure 1, Figure S2). These results once again emphasize the species-specific nature of the degree of utility of any new variable. Topographic variables are likely to be particularly useful for exploring competing hypotheses regarding whether local- to regional-scale factors such as microsite availability are important in determining species’ distributions (e.g., Bemmels et al. 2016).

Beyond general considerations about whether or not topographic variables are important for modeling a species’ distribution, care should also be taken in assessing whether or not static variables (i.e., variables that do not change over time) are appropriate to use for a given SDM application. The topographic variables we derive can be assumed to be largely static through time (especially in unglaciated regions, with the exception of changes in coastline reflecting sea-level changes). Stanton et al. (2012) explored the inclusion of static variables in SDM and found that including such variables when projecting to future climate-change scenarios typically improved, and rarely hindered, SDM performance when the variables were known to influence species distributions. Nonetheless, we recommend particular caution when projecting to contexts where topography may have changed substantially over the timescale of interest, for example due to Pleistocene glacial erosion in North America (Bell and Laine 1985).

### Conclusions

The envirem variables constitute a valuable dataset for species distribution modeling for a variety of applications. Although they are complementary to and largely derived from the WorldClim database that is already widely in use, they contain novel information not captured by this database. In particular, the envirem variables include conceptually novel climatic variables that may more closely reflect specific ecological and physiological processes, as well as topographic variables distinct from elevation that may represent non-climatic local- to regional-scale aspects of a species’ niche. In our exploration of case studies for 20 North American vertebrate species, the impact of including the envirem variables was species-specific: in 17 out of 20 cases model performance substantially improved compared to a model using only WorldClim variables, particularly when topographic envirem variables were included; in only three cases model performance was not substantially different or declined. In general, models built with and without the envirem variables produced habitat suitability predictions differing only modestly and at local scales in the current time period, but sometimes resulted in dramatic regional-scale differences in predicted habitat suitability when projected to a different time period. Overall, our results highlight how the envirem variables often improve model performance, even when biological information about the variables that are most relevant to determining habitat suitability for a given species is not known *a priori*. Furthermore, when knowledge about the determinants of species distributions is available from ecological theory, the envirem variables may be particularly useful for developing and testing the predictions of species-specific hypotheses. The significant improvements in model performance we observed for many species when following best practices in species distribution modeling suggest that the ENVIREM variables are worth general consideration for SDM, as their main benefit is providing a more comprehensive set of environmental variables to choose from, whether through statistical variable thinning or variable selection informed by ecological knowledge.

## Acknowledgements

We would like to thank L.L. Knowles for her guidance in developing this project. This manuscript greatly benefited from comments from G. Costa and anonymous reviewers. Funding was provided for graduate student support by an NSF GRFP fellowship (J.B.B.) and a University of Michigan Department of Ecology and Evolutionary Biology Edwin H. Edwards Scholarship in Biology (J.B.B.).

## Data accessibility

The envirem dataset has been deposited through the University of Michigan Deep Blue Data repository (DOI: XXXX), and can be accessed through the project website at envirem.github.io. The “envirem” R package is available on CRAN.

